# Generating human AMN and cALD iPSC-derived astrocytes with potential for modeling X-linked adrenoleukodystrophy phenotypes

**DOI:** 10.1101/2024.05.31.596696

**Authors:** Navtej Kaur, Jaspreet Singh

## Abstract

X-adrenoleukodystrophy (X-ALD) is a peroxisomal metabolic disorder caused by mutations in the ABCD1 gene encoding the peroxisomal ABC transporter adrenoleukodystrophy protein (ALDP). Similar mutations in ABCD1 may result in a spectrum of phenotypes in males with slow progressing adrenomyeloneuropathy (AMN) and fatal cerebral adrenoleukodystrophy (cALD) dominating the majority of cases. Mouse model of X-ALD does not capture the phenotype differences and an appropriate model to investigate mechanism of disease onset and progress remains a critical need. Induced pluripotent stem cell (iPSC)-derived and cell models derived from them have provided useful tools for investigating cell-type specific disease mechanisms. Here, we generated induced pluripotent stem cell (iPSC) lines from skin fibroblasts of two each of apparently healthy control, AMN and cALD patients with non-integrating mRNA-based reprogramming. iPSC lines expanded normally and expressed pluripotency markers Oct4, SOX2, Nanog, SSEA and TRA-1-60. Expression of markers SOX17, brachyury, Desmin, Oxt2 and beta tubulin III demonstrated the ability of the iPSCs to differentiate into all three germ layers. iPSC-derived lines from CTL, AMN and cALD male patients were differentiated into astrocytes. Differentiated AMN and cALD astrocytes lacked ABCD1 expression and accumulated VLCFA, a hallmark of X-ALD. These patient astrocytes provide disease-relevant tools to investigate mechanism of differential neuroinflammatory response and metabolic reprogramming in X-ALD. Further these patient-derived human astrocyte cell models will be valuable for testing new therapeutics.

## Introduction

The most common peroxisomal disorder affecting males at early ages, adrenoleukodystrophy (ALD), results from deletion/mutation in the ABCD1 gene, leading to an absent or non-functioning adrenoleukodystrophy protein (ALDP) [1, 2]. This defect causes an accumulation of very long chain fatty acids (VLCFA) in tissues and plasma via inhibition of peroxisomal β-oxidation [1, 3, 4]. The accumulation of VLCFA is the hallmark of X-ALD disease. Two major clinical variants exist: cerebral ALD (cALD) and adrenomyeloneuropathy (AMN). Though caused by the same or a similar mutation or deletion, cALD is biochemically associated with redox alterations, inflammation, and subsequent loss of myelin/oligodendrocytes [4]. cALD is often fatal in childhood, whereas AMN patients live to adulthood with mild involvement of the peripheral nervous system [1, 4-6].

Unlike human cALD, Abcd1-knock out (KO) mice do not show cerebral pathology [7-9]. However, they present AMN like symptoms and show imbalance in antioxidant systems with increasing age [10, 11]. The mechanisms that cause the spontaneous progression of disease from relatively mild AMN to fatal cALD remain unclear. We and others have documented a role for Abcd1-silenced astrocytes in X-ALD neuroinflammatory response [12-14]. Mice astrocytes silenced for Abcd1 and Abcd2 produced spontaneous inflammatory phenotype [13]. A similar inflammatory response was documented by our laboratory in Abcd1-KO mice primary astrocytes silenced for AMPKα1 [14]. Mouse astrocytes however could not be used to model differential inflammatory response seen in human AMN and cALD phenotypes.

Induced pluripotent stem cells (iPSCs) have become an attractive tool for in vitro disease modeling since they can give rise to any cell of the body. Skin fibroblasts were the first human cells to be reprogrammed to iPSCs due to ease of availability and growth in culture [15, 16]. In patient astrocytes obtained by directed differentiation iPSCs derived from skin fibroblasts of apparently healthy control (CTL), AMN and cALD male each we documented differential mitochondrial dysfunction, oxidative stress response, and neuroinflammatory cytokine profile [17]. In this work we generated iPSC-derived astrocyte from two additional male patients each of CTL, AMN and cALD phenotype. iPSC cells were in turn obtained by reprogramming of human skin fibroblasts from CTL, AMN and cALD.

## Results

### CTL, AMN and cALD patient fibroblast-derived iPSC are positive for AP and expressed pluripotency markers

CTL AMN and cALD human dermal fibroblasts were transfected with non-modified RNA (NM-RNA) containing, Oct4, Sox2, Klf4, cMyc, Nanog and Lin28 reprogramming factors and NM-miRNA as per the manufacturers protocol (ReproCell technologies). Daily overnight transfections were performed for four consecutive days and cells cultured in a hypoxic environment with 5% oxygen. Earliest emergence of iPSC colonies was observed ten days following the last transfection. Colonies had typical round and flat iPSC colony structure with a defined colony boundary separating it from the fibroblasts (Figure 1A). Pluripotency was established by checking for alkaline phosphatase activity (Figure 1B) and alkaline phosphatase live staining (Supplementary figure 1).

**Fig 1.**
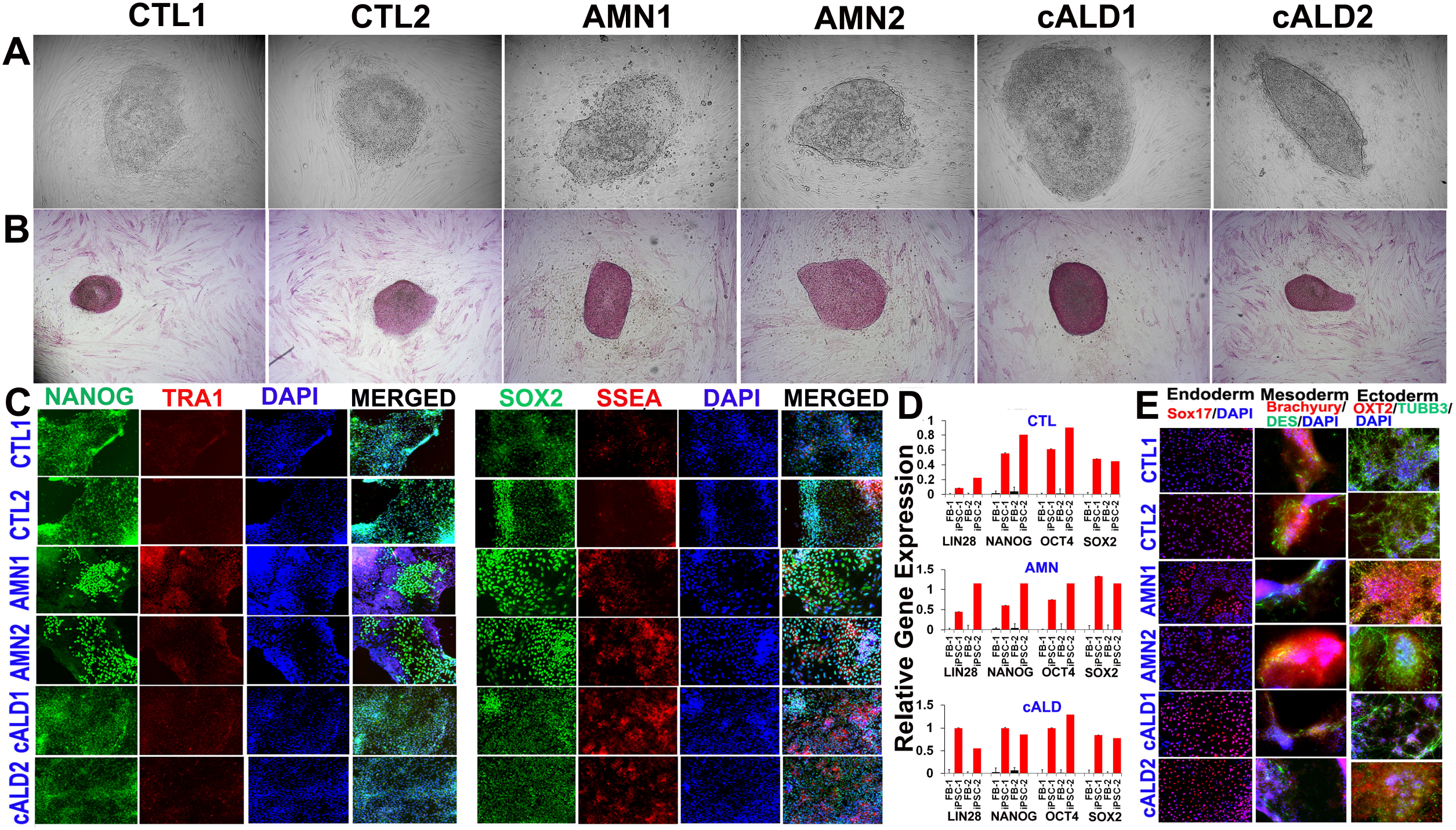

### Expression of pluripotency markers in CTL, AMN and cALD fibroblast-derived iPSCs

iPSC colonies were characterized for pluripotency markers after five passages. Fluorescent staining for pluripotent markers documented significant expression of NANOG, TRA-1-60, SOX2 and SSEA in iPSC colonies from CTL, AMN and cALD (Fig. 1C). RT-qPCR evaluation of CTL, AMN and cALD patient fibroblasts and the corresponding iPSC cells showed significantly higher expression of LIN28, NANOG, OCT4 and SOX2 in iPSC cells compared to fibroblasts (Fig. 1D).

### Differentiation of iPSC colonies to germ layers

Functional activity of iPSCs were determined by evaluating their ability to differentiate into three germ layers (Human pluripotent stem cell functional identification kit, R&D systems, Cat#SC027B). Immunofluorescent analysis of CTL, AMN and cALD cells disaggregated form embryoid bodies revealed various cell derivatives expressing ectodermal (OXT2 and β-III-tubulin), mesodermal (brachyury and desmin), and endodermal (Sox17) markers (Fig. 1E). Thus the obtained human iPSCs possess a broad differentiation potential *in vitro*.

### Astrocyte differentiation of CTL, AMN and cALD iPSC colonies

Monolayers cells were plated on Matrigel-coated plates in neural induction medium with SMAD inhibitor (SMADi) for 12-15 days to generate neural precursor cells using STEMdiff™ SMADi Neural Induction kit (STEMCELL Technologies). SMADi inhibits the expression of pluripotency genes and also suppresses differentiation in the mesodermal and ectodermal directions. Neural precursors were driven to astrocytic lineage by further culturing the cells on Matrigel-coated plates in astrocyte differentiation media for 18-20 days with media change every 2-3 days and passage very 6-8 days (STEMdiff™ Astrocyte differentiation kit, Stemcell Technologies). After 20-21 days on astrocyte differentiation medium, cells were incubated further in astrocyte maturation medium for 25-30 days with media changes every 2-3 days (STEMdiff™ Astrocyte maturation kit. STEMCELL Technologies). The cells were passaged every 6-8 days. After atleast three passages mature astrocytes were identified by immunofluorescence staining for mature glial cell markers including GFAP, aquaporin 4, S100β, ALDH1L1 and EAAT1 (Fig. 2A). Cells were negative for A2B5. ABCD1 mutation did not seem to affect the differentiation potential of iPSCs to astrocytes.

**Fig 2.**
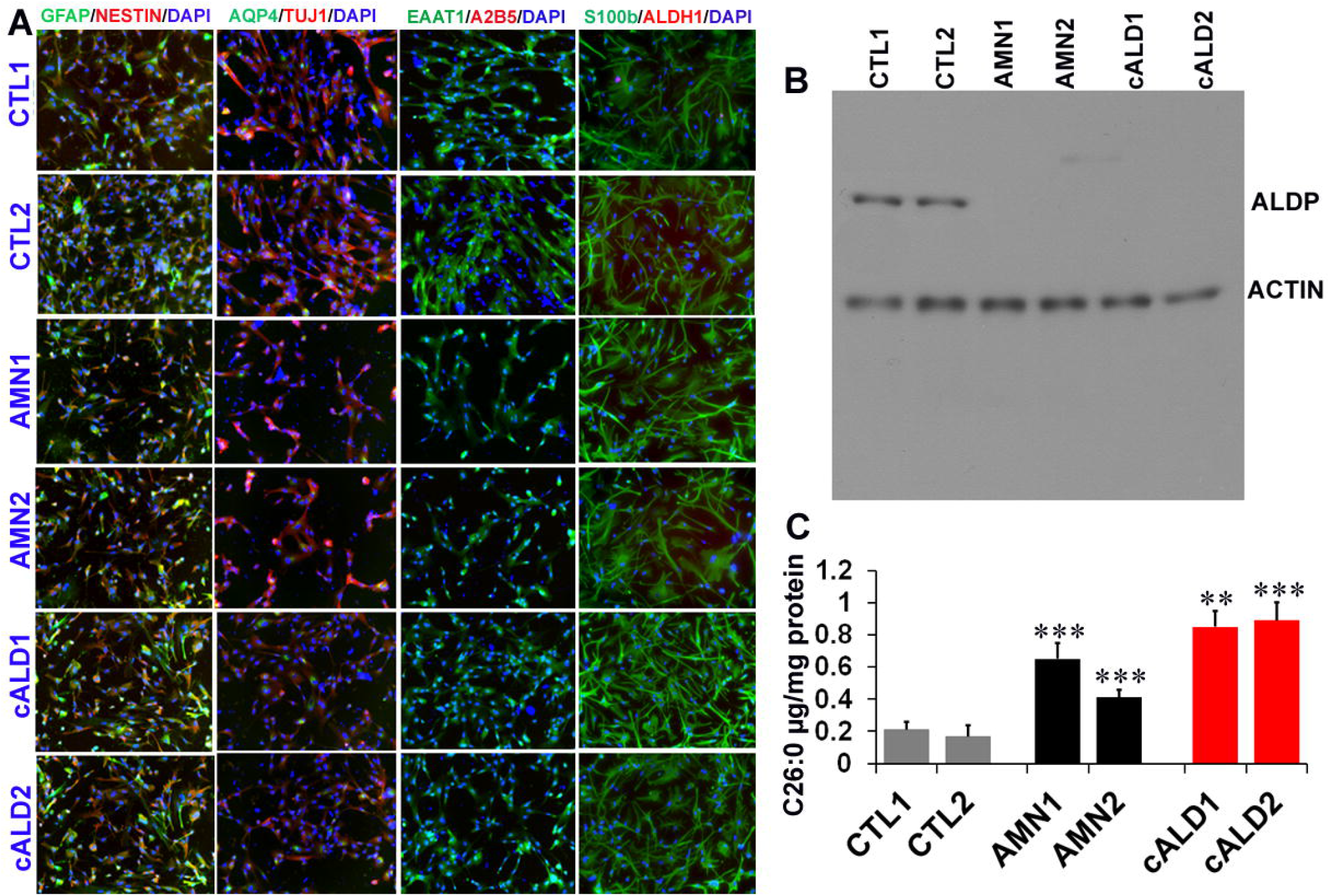

### ABCD1 expression and VLCFA levels in CTL, AMN and cALD iPSC-derived astrocytes

ABCD1 mutation is associated with loss of ALDP protein in AMN and cALD. We next investigated the status of ALDP in iPSC-derived CTL, Amn and cALD astrocytes by Western blotting. AMN and cALD iPSC-derived astrocytes exhibited loss of ALDP protein (Fig. 2B). Loss of ALDP is associated decreased β-oxidation and resultant VLCFA accumulation in X-ALD. We measured VLCFA (C26:0) levels in control, AMN and cALD iPSC-derived astrocytes. Absolute levels of C26:0 were significantly increased in AMN astrocytes compared to CTL astrocytes (Fig. 2C). The C26:0 levels in cALD were further increased significantly compared to AMN astrocytes (Fig. 2C).

### Mycoplasma detection in CTL, AMN and cALD iPSC-derived cells

Quality control of iPSC-derived cells were performed by nucleic acid amplification to detect mycoplasma species. No mycoplasma contamination was detected by PCR in any of the six CTL, AMN and cALD iPSC lines (Supplementary Figure 2).

## Discussion

X-ALD is the most frequent peroxisomal disorder affecting males, yet the disease mechanisms beyond ABCD1 mutation and VLCFA derangement largely remain unknown and there are no satisfactory therapeutic options. A significant barrier in unravelling the disease mechanisms has been lack of a relevant mouse model of the disease. X-ALD mouse model is a classical knockout of Abcd1 gene and accumulated VLCFA similar to human phenotype. It however, fails to develop CNS demyelination characteristic of the human cALD phenotype. Human iPSC-derived cell models of brain cells provide an opportunity to investigate early events in disease development and also to test novel therapeutic strategies. This study documents iPSC-derived astrocytes generated for apparently healthy controls (CTL), AMN and cALD phenotypes of X-ALD and adds biological replicates to our recent report [17]. Non-integrating mRNA-based reprogramming used in the present work is advantageous to generate clinical grade stem cells as there is no genome integration of the reprogramming factors. The non-modified mRNA and microRNA technology used to deliver the reprogramming factors (SOX2, LIN28, Oct4, KLF4, Nanog and cMyc) and immune evasion factors allow repeated transfections.

Documented role of astrocyte involvement in X-ALD neuropathology has been provided from human postmortem brain tissue and *in vitro* mouse astrocyte cultures by our laboratory and others [13, 14, 18-21]. Understanding the differential response of astrocytes in AMN and cALD phenotypes has been hampered by the lack of availability human primary cultures. This has started to change recently with the generation of iPSC-derived brain cell, including astrocytes from AMN and cALD phenotypes [17, 22, 23]. We recently documented generating astrocytes from iPSC-derived skin fibroblasts of one male patient each of CTL, AMN and cALD phenotype [17]. In the present study, we generated and characterized astrocytes obtained from iPSC lines reprogrammed from skin fibroblasts of additional two male patients each of CTL, AMN and cALD phenotype. In line with our recent report [17], ABCD1 mutation did not affect the reprogramming ability of AMN and cALD skin fibroblasts and the iPSC-lines were successfully differentiated into astrocytes. Immunocytochemical expression analysis of differentiated astrocytes indicated that they were positive for mature astrocyte markers including GFAP, EAAT, S100β and ALD1H1 consistent with previous studies [17, 22].

Mutations in ABCD1 gene encoding the ALDP protein is the primary clinical cause of X-ALD regardless of the phenotype [1]. We observed that similar to AMN and cALD patient fibroblasts, the ABCD1 expression was lacking in the AMN and cALD iPSC-derived astrocytes. Lack of ALDP is associated with accumulation of VLCFA in patient body fluids and tissues including the central nervous system [4]. While plasma VLCFA levels do not correlate with disease phenotype variability in male patients, VLCFA is still the only biochemical hallmark of the disease [3, 4, 6, 24]. Mouse astrocytes silenced for Abcd1 and Abcd2 and human transformed cell lines silenced for Abcd1 also accumulate VLCFA [13, 21, 25]. However, they are unable to account for the differential response seen in AMN and cALD phenotype. We recently documented differential accumulation of VLCFA in patient iPSC-derived astrocytes from AMN and cALD phenotypes [17]. In line with these, VLCFA levels were also differentially increased in AMN and cALD patient iPSC-derived astrocytes in the present study with higher accumulation in cALD patient iPSC-derived astrocytes. These results also support previous studies of higher accumulation of VLCFA in cALD patient iPSC-derived brain cells [22, 23].

Combining the iPSC-derived astrocytes from the present study with our recent report, we have multiple patient iPSC-derived brain cells from AMN and cALD phenotypes and age- and sex-matched controls. These patient-derived cells can be a useful tool for biomarker discovery in addition to the mechanistic studies. We recently documented the role of microRNAs (miRNAs) and metabolites in CNS disease process in cALD human postmortem brain tissue and as plasma biomarkers of disease severity in AMN [26, 27]. Human iPSC-derived astrocytes developed in this study and our recent report can help decipher the role played by miRNAs and metabolites in astrocyte-mediated neuroinflammation in early stages of the disease and progression in X-ALD phenotypes.

## Material and Methods

### Ethics Approval

The study protocol was approved by institutional IRB (#13352). Fibroblast samples were de-identified specimens obtained from Coriell Cell repositories and did not involve recruitment of human subjects.

### Human fibroblasts

Human skin fibroblast of healthy CTL fibroblasts (GM08402; 32-year old male, GM03348; 10-year old male), AMN fibroblasts (GM17819; 32 year old male, GM07675; 22 year old male), and cALD (GM04904; 11-yearold male, GM04496; 6-year old male) were obtained from the National Institute for General Medical Sciences human genetic cell repository at Coriell Institute for Medical Research, Camden, NJ.

### Derivation of iPSCs and Differentiation into Astrocytes

iPSCs were generated from skin fibroblasts with the Stemgent® StemRNA™ 3rd Gen Reprogramming Kit (Reprocell). The iPSCs were functionally characterized by checking for their ability to differentiate into the three germ layers (Human pluripotent stem cell functional identification kit, R&D systems, Cat#SC027B) according to the manufactures protocol. Neural Precursor Cells (NSCs) were generated from human induced pluripotent stem cells (iPSCs) using STEMdiff™ SMADi Neural Induction kit (STEMCELL Technologies) via dual SMAD inhibition as per the manufacturers protocol. The NPC were differentiated with STEMdiff™ Astrocyte differentiation kit and STEMdiff™ Astrocyte maturation kit (STEMCELL Technologies) as per the manufacturers protocol.

### Mycoplasma detection

Presence of mycoplasma was checked by PCR using mycoplasma detection kit (Venor^®^ GeM OneStep, Minerva biolabs).

### Culturing of mature iPSC-derived astrocytes

Astrocytes were cultured at seeding density of 5 x 10^4^ cells/cm^2^ (day 0) at 37°C and 5% CO_2_ in in astrocyte medium (Neurobasal-A medium, Thermo Fisher Scientific) containing N21 max (1X, R&D Systems), FBS One Shot (1X, Thermo Fisher Scientific), Glutamax (1X, Gibco, Thermo Fisher Scientific), heregulin-β1(10 ng/mL, Peprotech, Inc.), bFGF (8ng/mL, R&D Systems), penicillin-streptomycin (1X, ThermoFisher Scientific).

### VLCFA Analysis

CTL, AMN, and cALD astrocytes (2.5 x 10^5^ cells each) were processed at Wayne State University Lipidomics Core facility. Saturated (hexacosanoic [C26:0] was calculated as per microgram of protein. Lipids were subject to alkaline methanolysis and resulting fatty acid methyl esters were analyzed by gas chromatography-mass spectrophotometry (QP2010 GC-MS system, Shimadzu Scientific Instruments) equipped with Restek column, as reported [5].

### Quantitative Real-Time Polymerase Chain Reaction Gene Expression

Total RNA was extracted with the miRNeasy kit (Qiagen) and 1µ RNA was used for cDNA synthesis. RT-qPCR were conducted using CFX96 Real-Time PCR Detection System (BioRad) using IQ SYBR Green Supermix (BioRad), as described previously [17]. Gene expression was normalized to 60S ribosomal L27 gene and samples were run in triplicate. Primer sequences of genes investigated are listed in the Appendix 1.

### Immunofluorescence Staining

**CTL**, AMN and cALD cells were plated on Matrigel coated chambered slides (25000 cells/cm^2^) and allowed to grow overnight. Cells were fixed with 4 % paraformaldehyde for 15 mins at room temperature (RT), washed with PBS, and incubated in blocking solution containing 10 % normal donkey serum (Sigma) and 0.03 % Triton X-100 (Sigma-Aldrich) in PBS for 1 hr at RT. Primary antibodies were incubated overnight at 4 °C and secondary antibodies for 1 hr at RT. Nuclei were counterstained with DAPI (Sigma). Images were acquired with Fluorescence microscope (BZ-X series, Keyence). Antibodies used are listed in the Appendix 2.

### Western Blot Analysis

CTL, AMN and cALD astrocytes were homogenized in radioimmunoprecipitation (RIPA) buffer with protease inhibitor cocktail (Thermo Fisher Scientific). 60 μg of total protein was electrophoresed as described previously [17]. Antibodies used are listed in the Appendix 2.

## Data Analysis

Data was analyzed using GraphPad Prism software (version 7.0). Normality was assessed with Kolmogorov-Smirnov test. Groups were compared with two-tailed unpaired Student’s t-test for normally distributed or nonparametric Mann-Whitney test for non-normally distributed data.

Statistical significance was set at *p*<0.05.

## Authorship contribution statement

JS conceived the idea, designed the experiments, provided direction and funding. NK designed and performed the experiments, contributed to the acquisition and analyses of data and drafted the manuscript. All authors revised and approved the final version of the manuscript.

