## Appendix 1 for "Generating human AMN and cALD iPSC-derived astrocytes with potential for modeling X-linked adrenoleukodystrophy phenotypes"

**Human Real time PCR primer sequences:**

Oct4-F: GAAACCCACACTGCAGATCA
Oct4-R: GGTTACAGAACCACACTCG

Nanog-F: AGATGCCTCACACGGAGACT
Nanog-R: TTTGCGACACTCTTCTCTGC

Sox2-F: TGCTGCCTCTTTAAGACTAGGAC
Sox2-R: CCTGGGGCTCAAACTTCTCT

Lin28-F: GGCAGTGGAGTTCACCTTTAAGA
Lin28-R: AGCTTGCATTCCTTGGCATGATGA

L27-F: TGGACAAAACTGTCGTCAATAAGG
L27-R: AGAACCACTTGTTCTTGCCTGTC
