## Appendix 2 for "Generating human AMN and cALD iPSC-derived astrocytes with potential for modeling X-linked adrenoleukodystrophy phenotypes"

| Catalog No | Antibody Details |
|---|---|
| 5685S | EAAT1 (D20D5) RABBIT mAb |
| 59678S | AQP4(D1F8E) XP RABBIT mAb |
| 85828S | ALDH1L1(E712Q) RABBIT mAb |
| G3893 | MONOCLONAL ANTI-GLIAL FIBRILLARY ACID PROTEIN (GFAP) |
| S2532 | MONOCLONAL ANTI-S-100B |
| AB53521-1001 | Anti-A2B5 antibody |
| 5568S | BETA-3-TUBULIN |
| 5332S | Desmin (D93F5) XP® Rabbit mAb |
| 963121 | Goat anti-human SOX17 |
| 963273 | Goat anti-human Otx2 |
| 963427 | Goat anti-human Brachyury |
| 73349S | NESTIN |
| AF2018 | Human/Mouse/Rat SOX2 Affinity Purified Polyclonal Ab |
| 60064AD | Anti-Human TRA-1-60 Antibody, Clone TRA-1-60R, Alexa Fluor® 488 |
| 60062PE | Anti-Human SSEA-4 Antibody, Clone MC-813-70, PE |
| 2840S | Oct-4A Rabbit mAb |
| AF1997 | Human Nanog Antibody |
| MAB2018 | Human/mouse/rat Sox2 |
| 2840S | Oct-4A Rabbit mAb |
| AF-1759 | Human/Mouse/Oct-3/4 Antibody |
| 23064S | Sox2 (D9B8N) Rabbit mAb |
| ab197013 | ABCD1/ALD antibody [EPR15929] |

| **Secondar Antibodies** | |
|---|---|
| A21206 | Alexa Fluor 488 Donkey Anti Rabbit |
| A21042 | Alexa Fluor 488 Goat anti mouse |
| A11055 | Alexa Fluor 488 Donkey anti goat |
| 705-605-147 | Alexa Fluor 647 Donkey Anti Goat |
| 715-605-150 | Alexa Fluor 647 Donkey Anti mouse |
| 711-605-152 | Alexa Fluor 647 Donkey Anti Rabbit |

| **Manufacturer** | **Dilution** |
|---|---|
| CELL SIGNALING | 1/200 |
| CELL SIGNALING | 1/400 |
| CELL SIGNALING | 2/100 |
| SIGMA | 1/100 |
| SIGMA | 1/100 |
| ABCAM | 1/100 |
| Cell Signaling | 1/200 |
| Cell Signaling | 1/100 |
| R&D | 1/100 |
| R&D | 1/100 |
| R&D | 1/100 |
| CELL SIGNALING | 1/400 |
| R&D | 5 µg/ml |
| STEM CELL TECH | 2/100 |
| STEM CELL TECH | 2/100 |
| Cell Signaling | 1/100 |
| R&D | 10 µg/ml |
| R&D | 8 µg/ml |
| Cell Signaling | 1/100 |
| R&D | 10 µg/ml |
| Cell Signaling | 1/100 |
| Abcam | 1/1000 |

| | |
|---|---|
| Invitrogen | 1/200 |
| Invitrogen | 1/200 |
| Invitrogen | 1/200 |
| JacksonImmunoResearch | 1.5/100 |
| JacksonImmunoResearch | 1.5/100 |
| JacksonImmunoResearch | 1.5/100 |
