## Supplementary figure for "Generating human AMN and cALD iPSC-derived astrocytes with potential for modeling X-linked adrenoleukodystrophy phenotypes"

SUPPLEMENTARY FIGURES:

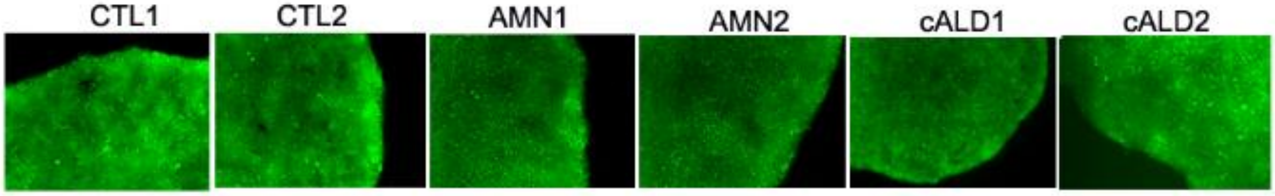


**Supplementary Figure S1**: Alkaline phosphatase live staining. Magnification 10X

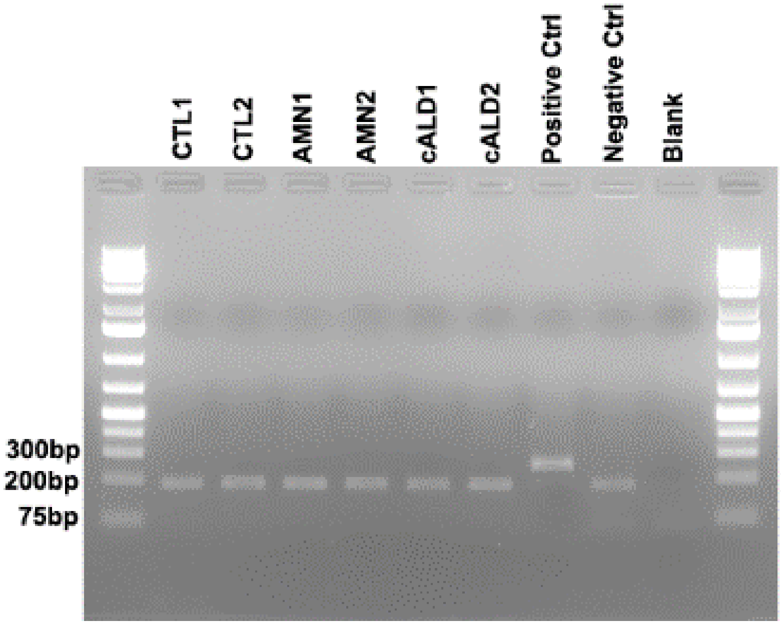


**Figure S2:** Mycoplasma detection
